# *Enterococcus* spp. have higher fitness for survival, in a pH-dependent manner, in pancreatic juice among duodenal bacterial flora

**DOI:** 10.1101/2021.08.08.455373

**Authors:** Saki Itoyama, Emika Noda, Shinji Takamatsu, Jumpei Kondo, Rui Kawaguchi, Munefumi Shimosaka, Tomoya Fukuoka, Daisuke Motooka, Shota Nakamura, Masahiro Tanemura, Suguru Mitsufuji, Yoshifumi Iwagami, Hirofumi Akita, Toru Tobe, Yoshihiro Kamada, Hidetoshi Eguchi, Eiji Miyoshi

## Abstract

**Objectives:** Bacterial infection is involved in the progression of many gastrointestinal diseases, including cancer; however, how and which bacteria colonize in pancreatic juice and tissue have yet to be elucidated. Recently, we reported that *Enterococcus faecalis* exists in the pancreatic juice and tissues of patients with chronic pancreatic disease. Here, we investigated the survival of *E. faecalis* in duodenal juice with different pH conditions.

**Methods:** Pancreatic juice samples from 62 patients with cancers of the duodeno-pancreato-biliary region were evaluated for the presence of *E. faecalis*. 16S ribosomal RNA PCR and 16S-based metagenome analyses were performed to determine the bacterial composition. The survival of *E. faecalis* in various pancreatic juice conditions was evaluated.

**Results:** Of 62 samples, 27% (17/62) were positive for *Enterococcus* spp., among which 71% (12/17) contained *E. faecalis. Enterococcus* spp. showed the highest fitness for survival in alkaline pancreatic juice among various bacterial species. The microbiome of pancreatic juice from patients with pancreatic and bile duct cancer showed diversity, but *Enterococcus* spp. were enriched among duodenal tumors and intraductal papillary mucinous neoplasms.

**Conclusions:** Alkalinity is important for the selective survival of *E. faecalis* among microbiota. *E. faecalis* may induce pancreatic inflammation with changes in pancreatic juice conditions.

## Introduction

Pancreatic juice, or pancreatic fluid, in healthy individuals is thought to be sterile due to its antibacterial activity toward a large spectrum of bacteria^1,2^. In a normal healthy situation, both the alkalinity of the fluid and small-molecule peptides in the fluid protect the pancreas against ascending bacterial infection from duodenal microbiota^2^. However, bacterial infection is present in the pancreatic juice of patients with chronic pancreatic disease, including chronic pancreatitis^3-6^ and pancreatic cancer^3,5,7^. Although bacterial infection in pancreatic juice has been regarded as a complication that affects the severity and therapeutic course of pancreatitis, the role of bacteria in pancreatic cancer development, carcinogenesis, and treatment is being investigated^8^.However, the mechanism by which bacteria become able to colonize in pancreatic juices, as well as the type of bacteria involved, has yet to be elucidated.

*Enterococcus faecalis* is a gram-positive commensal bacterium that is part of the normal gut microbiota in healthy individuals^9^. *E. faecalis* and *E. faecium*, the most abundant species of the *Enterococcus* genus in humans, have recently emerged as a cause of nosocomial infection, in particular urinary tract infections^10^. We previously demonstrated the presence of *Enterococcus* spp. in pancreatic juice obtained from patients with cancers of the duodeno-pancreato-biliary region. Additionally, we showed an increase in serum antibodies against *E. faecalis* capsular polysaccharide in patients with chronic pancreatitis and pancreatic cancer, as well as the possible involvement of *E. faecalis* in the progression of pancreatic diseases^5^. In this previous report, the presence of *E. faecalis* in pancreatic juice was evaluated using a genetic and immunohistological approach; however, we have not confirmed the colonization of live *E. faecalis* in pancreatic juice. Here, we isolated live *E. faecalis* from pancreatic juice samples of patients with duodeno-pancreato-biliary disease. Additionally, we evaluated the effect of pancreatic juice pH levels on the survival of *E. faecalis* in pancreatic juice from the duodenum.

## Materials and Methods

### Pancreatic juice and tissue

The ethical committee at Osaka University Hospital approved the study protocol (protocol IDs 14107 and 15212), and written informed consent was obtained from each participant. Pancreatic juice was collected after pancreatectomy from the drainage tube from 62 patients, including 34 patients with pancreatic cancer and 28 with duodenal or bile duct cancer. None of the patients underwent previous sphincterotomy There were no cases of distal pancreatectomy. Only clear pancreatic juice was used in this study. All samples were collected at Osaka University Hospital, National Hospital Organization Kure Medical Center or Osaka Police Hospital and kept frozen at −80°C or refrigerated at 4°C until use.

### Detection of bacterial DNA in pancreatic juice and bile by PCR analysis

Pancreatic juice was used directly as the template for polymerase chain reaction (PCR). PCR amplification was performed in 20-µl reactions containing forward and reverse primers specific for the bacterial 16S ribosomal RNA (rRNA) gene^11^. The primers used in this study and temperature profiles for PCR are listed in Supplemental Table 1.

### 16S ribosome metagenome analysis of bacteria in duodenal and pancreatic juices

16S rRNA metagenome analysis was performed at the Genome Information Research Center in Research Institute for Microbial Diseases, Osaka University. In brief, bacterial DNA was extracted from pancreatic juice or duodenal juice samples using a DNeasy PowerSoil Pro kit (Qiagen, Venlo, Netherlands). Each library was prepared according to the Illumina 16S Metagenomic Sequencing Library Preparation Guide with the following primer sets targeting the V1–V2 region of 16S rRNA genes:27Fmod, 5′-AGRGTTTGATCMTGGCTCAG-3′; and 338R, 5′-TGCTGCCTCCCGTAGGAGT-3′. For the amplicons, 251-bp paired-end sequencing was performed on a MiSeq system (Illumina, CA, USA) using a MiSeq Reagent v2 500 cycle kit. The paired-end sequences obtained were merged, filtered, and denoised using DADA2 (https://github.com/benjjneb/dada2). Taxonomic assignment was performed using the QIIME2 feature-classifier plugin with the Greengenes 13_8 database. The QIIME2 pipeline, version 2020.2, was used as the bioinformatics environment for the processing of all relevant raw sequencing data (https://qiime2.org).

### Bacterial culture and isolation of *E. faecalis*

To isolate live bacteria, pancreatic juice was cultured for 24 hours at 37°Ceither with brain-heart infusion (BHI) agar (3.7% BBL™ Brain Heart Infusion [BD, NJ,USA], 1.35% Bacto™ Agar [BD] in H_2_O) or Enterococcosel agar (5.6% BBL™Enterococcosel™ Agar [BD] in H_2_O) for the purpose of culturing entire bacterial spectra or selecting *Enterococcus* spp., respectively.

To isolate *E. faecalis*, colonies grown on Enterococcosel agar were isolated and inoculated in Colombia media (3.5% Difco™ Columbia broth [BD] in H_2_O). A portion of each bacterial culture was subjected for PCR analysis to detect *E. faecalis*–specific 16S rRNA using the specific primer set listed in Supplemental Table 1. Isolated bacterial cultures confirmed to be *E. faecalis* were frozen as glycerol stocks until further use.

### Survival of duodenal bacterial flora in pancreatic juice

Aseptic pancreatic juice was selected by PCR-based screening with a nonspecific 16S rRNA primer set (Supplemental Table 1). Duodenal juice was mixed with either aseptic pancreatic juice or sterile phosphate-buffered saline (PBS) at a ratio of 1:49 and incubated at 37°C for 24 hours. Then, part of the mixture was analyzed for the constituents by 16S rRNA metagenome analysis, as described above. A separate portion of the mixture was spread onto Enterococcosel agar and cultured for 24 hours at 37°C.

### Bacterial culture in various conditions of pancreatic juice

Colorimetric pH measurement was performed for 12 independent pancreatic juice samples by applying 1 µL sample onto MColorpHast™ pH strips (Merck Millipore, MA, USA). To adjust the pH of pancreatic juice, 4 µL NaHCO_3_ (1 M) was added to 46µL pancreatic juice. After the adjustment, colorimetric measurement was completed to confirm the pH was within the physiological range (9.0-9.5). Each pancreatic juice sample (50 µL) before and after pH adjustment was inoculated in Enterococcosel agar and incubated for 24 hours. After 24 hours cultivation, each plate was assigned to one of five groups according to the number of colonies.

To evaluate the fitness in various pancreatic juices, *E. faecalis* cultured in Columbia medium was adjusted to OD 690 of 1.0 and diluted to 1:10^−6^ in pancreatic juice from 3 different patients wherein no bacterium was detected. After incubating at 37°C overnight, 10 µL culture was spread onto BHI agar and further cultured for 24 hours.

### Statistical analyses

Statistical analyses were performed using either the Wilcoxon rank-sum test or Student’s *t*-test. A *P* value less than 0.05 was considered statistically significant.

## Results

### *Enterococcus* spp. are enriched in pancreatic juice among duodenal bacterial flora

To evaluate the ability of *Enterococcus* spp. to survive in pancreatic juice,duodenal juices containing bacterial flora were cultured in aseptic pancreatic juice samples for 24 hours. The enrichment of *Enterococcus* spp. was visualized by an expansion of colonies on Enterococcosel agar, an *Enterococcus* spp. selective agar plate (Fig. 1A). Additionally, 16S rRNA metagenome analysis confirmed the enrichment of *Enterococcus* spp. (Fig. 1B, C). These results suggest that *Enterococcus* spp. have a higher potential to survive in pancreatic juice among duodenal bacterial flora, which is the most likely source of bacterial colonization in bile and pancreatic juices.

**Figure 1.**
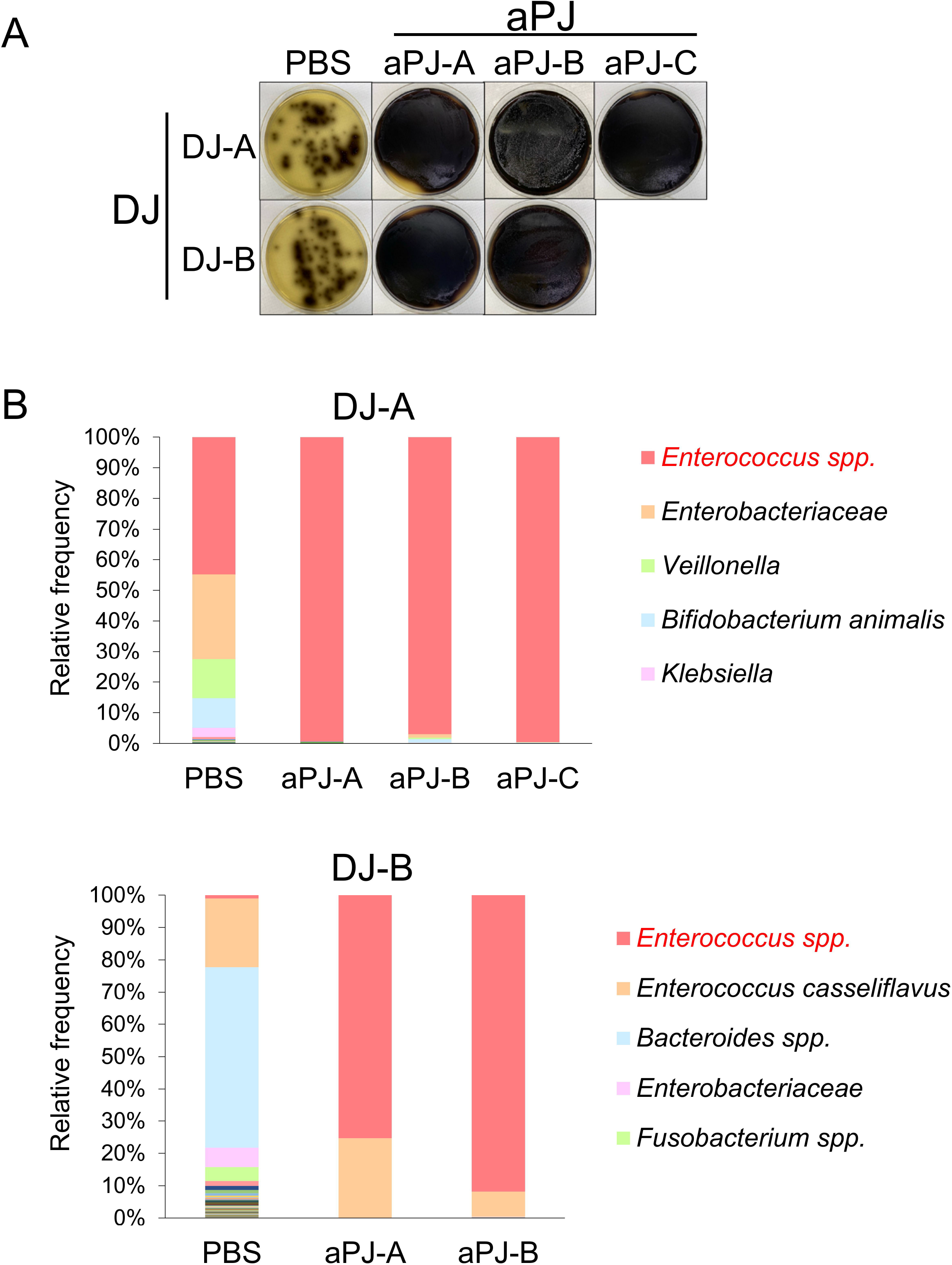
*Enterococcus* spp. have higher fitness to survive in pancreatic juice. (A) Visual agar plate assay to detect the abundance of *Enterococcus* spp. in a mixture of duodenal juice (DJ) and aseptic pancreatic juice (aPJ) at a 1:49 volume ratio. The DJ/aPJ mixture was incubated for 24 hours before plating on Enterococcosel agar. Phosphate-buffered saline was used as a control instead of aPJ. (B) Metagenome analysis of the DJ/aPJ mixture. After incubation, the samples were subjected to 16S rRNA metagenome analysis without selection on the agar plate. Each colored bar indicates the relative frequency of indicated bacterial species, which are listed on the right panel. aPJ-A, aseptic pancreatic juice from ampullary cancer; aPJ-B, aseptic pancreatic juice from pancreatic neuroendocrine tumor; aPJ-C, aseptic pancreatic juice from duodenal cancer

Healthy pancreatic juice is thought to be aseptic, and this is explained in part by its alkaline condition^2^. To evaluate the pH level of pancreatic juice in disease and its correlation with bacterial colonization, pancreatic juice samples from 62 patients were analyzed. Of 62 samples, 45% (28/62) were positive for bacterial culture, of which 61% (17/28) were positive for *Enterococcus* spp. (Fig. 2A). Among *Enterococcus* spp., 71% (12/17) contained *E. faecalis* (Fig. 2A). The pH of pancreatic juice varied from 7.5 to 9.5, with the pH of most samples lower than that of normal pancreatic juice (Fig. 2B-D).Interestingly, the pH of pancreatic juice in which bacteria were detected was significantly lower than that of aseptic pancreatic juice (Fig. 2B). Despite the correlation between lower pH and general bacterial colonization in pancreatic juice, the detection of *Enterococcus* spp. and *E. faecalis* was not significantly correlated with a lower pancreatic juice pH (Fig. 2C, D). These results support the potential of *Enterococcus* spp., including *E. faecalis*, to survive in pancreatic juice.

**Figure 2.**
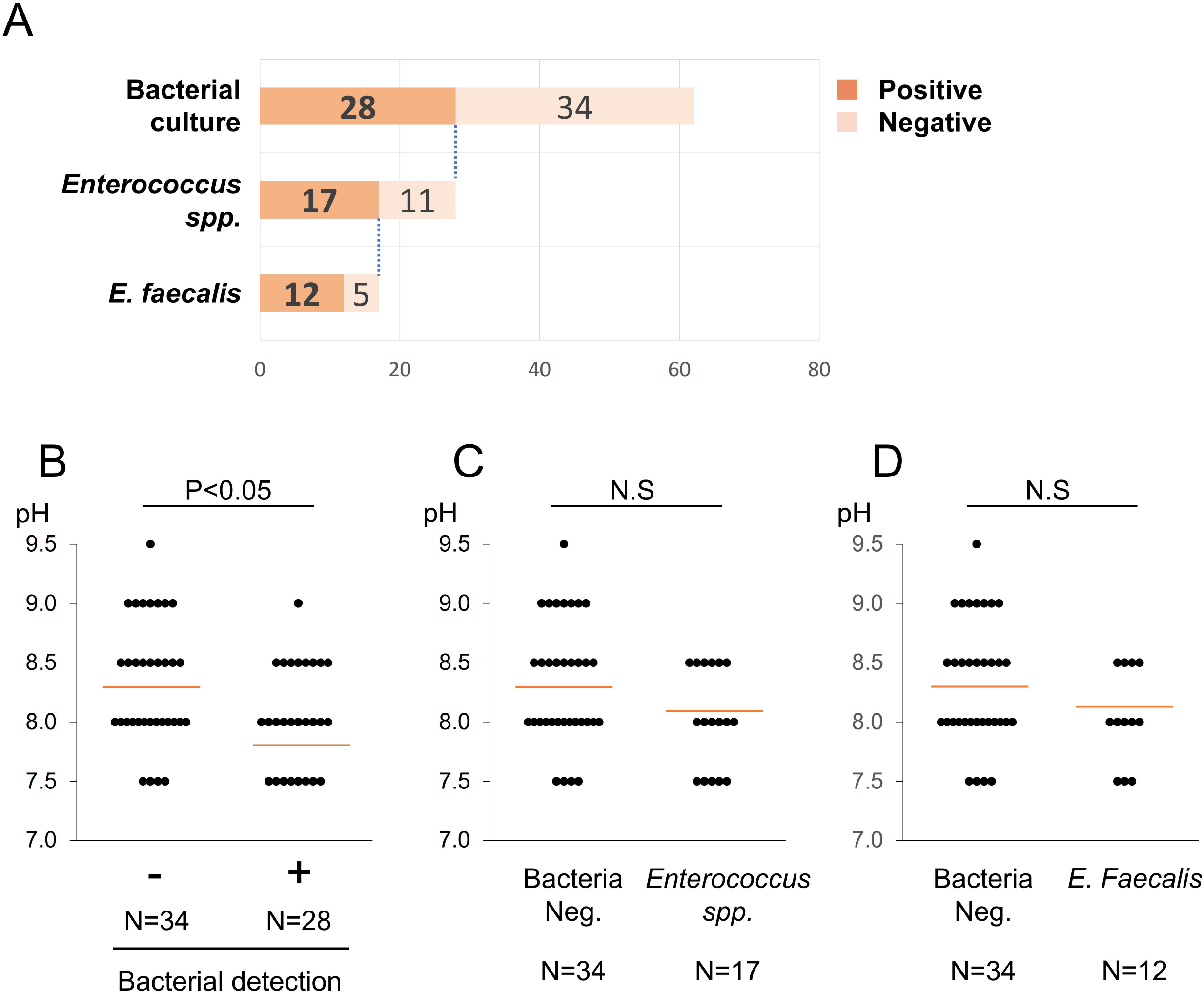
*Enterococcus* spp. can survive in pancreatic juice with a higher pH than other bacteria. (A) A chart showing the detection of *Enterococcus faecalis* from 62 pancreatic juice samples. (B-D) Comparison of the pH level in pancreatic juice samples with negative versus positive culture for any bacteria (B), *Enterococcus* spp. (C), and *E. faecalis* (D). The number of the samples assigned for each group is indicated below the bars. Red lines designate the average in each group. Wilcoxon rank sum test was used for the statistical analysis.

### *Enterococcus* spp., including *E. faecalis*, can colonize pancreatic juice at subphysiological pH levels

Although *Enterococcus* spp. have higher survival potential in pancreatic juice,bacteria were not detected in any of the pancreatic juice samples with a pH from 9 to 9.5 with only one exception. This is consistent with the common recognition that pancreatic juice is sterile in healthy patients due to its alkaline condition. This leads to the hypothesis that the slightly lowered pH of pancreatic juice, which can occur in pathogenic situations such as chronic pancreatitis, can be a trigger that allows colonization of *Enterococcus* spp., including *E. faecalis*.

To investigate this hypothesis, we evaluated the fitness of *E. faecalis* in pancreatic juice at different pH levels. *E. faecalis* strains isolated from 12 different pancreatic juice samples were cultured in pancreatic juice samples from pancreatic cancer patients with a pH of 9.5, 9.0, or 8.5, and colony formation capacity was measured. Our results showed that survival of all 12 strains was suppressed in pancreatic juice with a pH 9.5, whereas 5 in 12 and 11 in 11 strains survived in pancreatic juice with a pH 9.0 and 8.5, respectively (Fig. 3A, Supplemental Fig. S1). To further support the correlation between bacterial fitness in pancreatic juice and its pH,bacterial culture was completed for pancreatic juice samples with increased pH within the physiological range (pH, 9.0-9.5). pH levels were adjusted by the addition of NaHCO_3_ to the pancreatic juice samples. Consistent with the above results, colony formation was suppressed in 75% (9/12) of the pancreatic juice samples compared with the nonadjusted culture (Fig. 3B, Supplemental Fig. S2). These results indicate that *Enterococcus* spp. can survive pancreatic juice at subphysiological pH levels.

**Figure 3.**
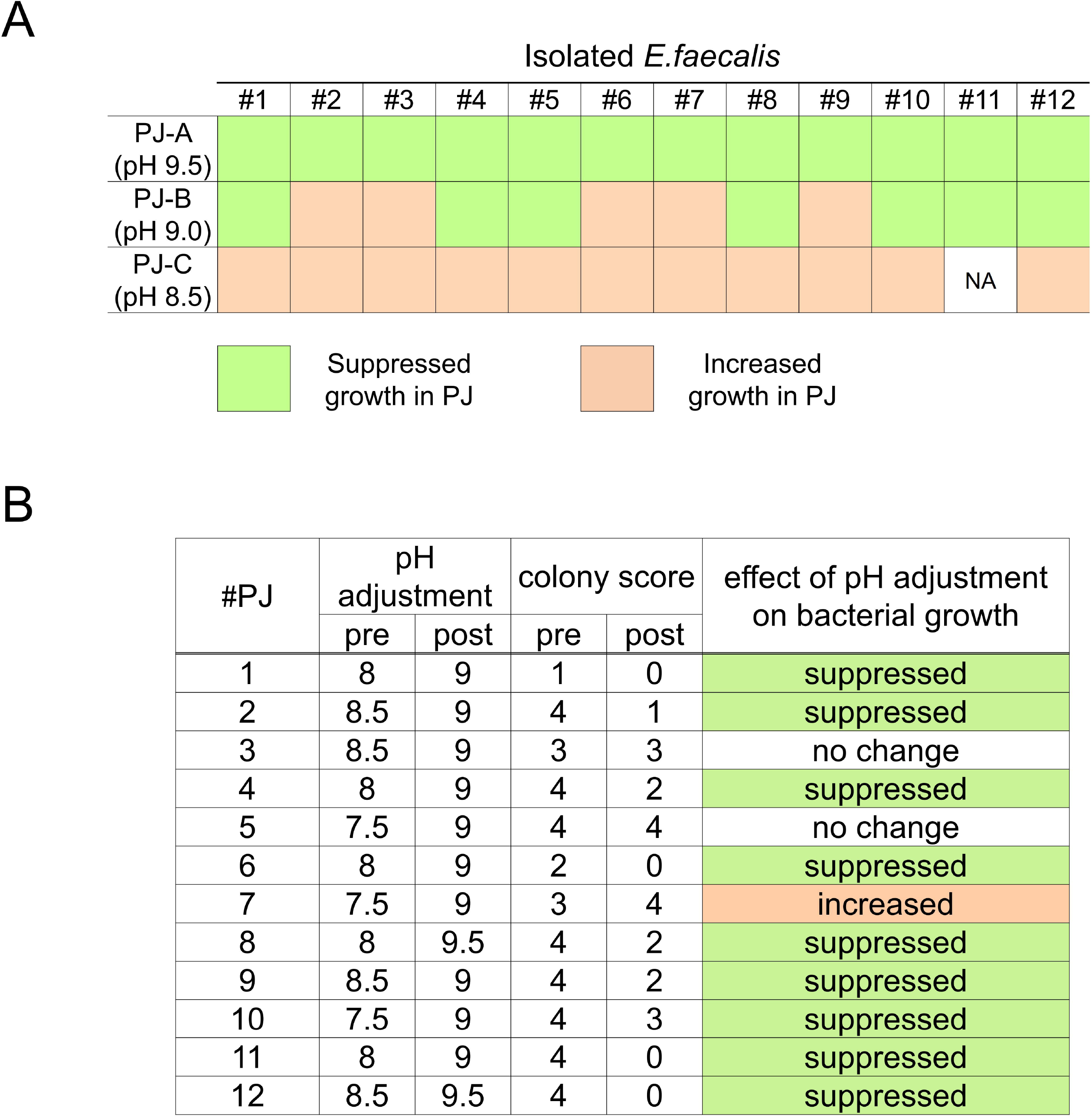
*Enterococcus* spp. shows suppressed growth in pancreatic juice with a pH above physiological limit. (A) Visual agar plate assay to detect the fitness of isolated *Enterococcus faecalis* to pancreatic juice with different pH levels. Twelve isolated *E. faecalis* samples were incubated with each pancreatic juice sample and plated on BHI agar. The changes in bacterial growth compared with the control (phosphate-buffered saline) were shown. See Supplemental Figure S1 for the original images of the agar plates. (B) Visual agar plate assay to detect the fitness of *Enterococcus* spp. in pancreatic juice. The 12 pancreatic juice samples, with or without adjusted pH levels, were plated on Enterococcosel agar. Bacterial growth was scored according to number of colonies, as shown in Supplemental Figure S2. NA, non-assessable

### *Enterococcus* spp. are enriched in the pancreatic juice of diseased pancreas with intact pancreatic and biliary ductal flow, but not with cancer in the pancreatobiliary region

Although *Enterococcus* spp. and *E. faecalis* were enriched in pancreatic juice samples compared with other bacterial species, bacterial flora was not enriched for *Enterococcus* spp. in the samples of pancreatic cancer patients (Supplemental Fig. S3) when compared with matched duodenal juice samples. To understand this discrepancy, we analyzed the relationship between the frequency of *Enterococcus* spp. and the disease type for each case. Metagenome analysis was performed on pancreatic juice samples from 11 patients who had undergone pancreatoduodenectomy: 5 for pancreatic or bile duct cancers, 2 for intraductal papillary mucinous neoplasia (IPMN), and 4 for duodenal tumors. The pancreatic juice samples from five pancreatobiliary cancer patients exhibited a limited frequency of *Enterococcus* spp., whereas those from IPMN and duodenal tumor patients exhibited clear enrichment of *Enterococcus* spp. (Fig. 4A). Interestingly, the pH of each group (5 samples from pancreatobiliary cancers vs. 6 samples from the others) was not statistically different (Fig. 4B; p > 0.05 by Wilcoxon rank sum test), indicating the presence of factors in addition to the pH level allowing bacterial species other than *Enterococcus* spp. to colonize pancreatic juice.

**Figure 4.**
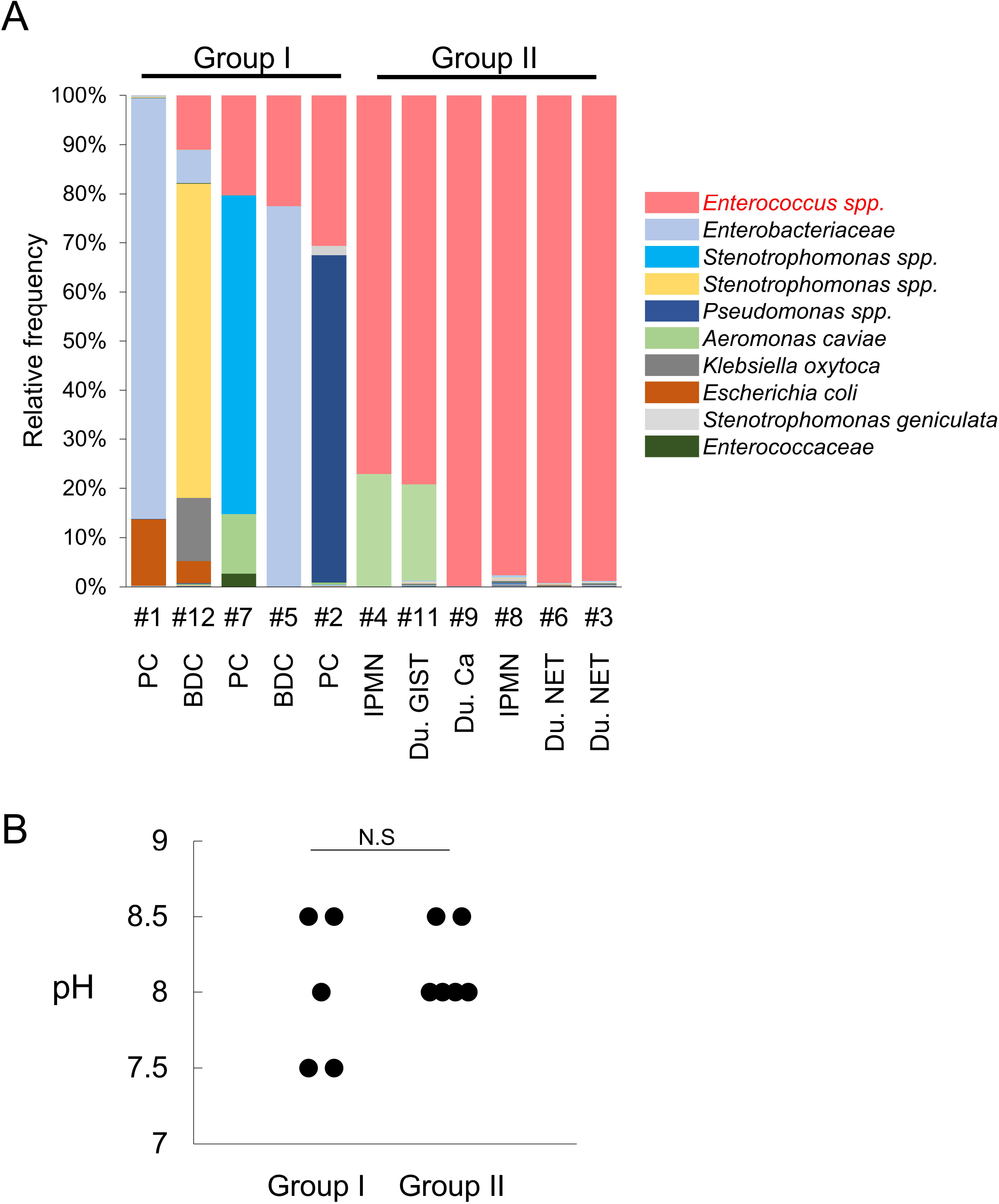
Microbiota in pancreatic juice is diverse in patients with pancreatobiliary cancer but is highly enriched with *Enterococcus* spp. in patients with duodenal region tumors and IPMN. (A) Metagenome analysis of 11 pancreatic juice samples. 16S rRNA metagenome analysis was performed on pancreatic juice samples from the indicated cases. Each colored bar indicates the relative frequency of the indicated bacterial species, which are listed on the right panel. BDC, bile duct cancer; Du, duodenal-; IPMN, intraductal papillary mucinous neoplasia; PC, pancreatic cancer. (B) Comparison of pH levels in pancreatic juice samples of patients with pancreatobiliary cancer versus those with duodenal tumors and IPMN. Wilcoxon rank sum test was used for the statistical analysis.

## Discussion

In the present study, we showed *Enterococcus* spp. have a higher potential to survive and colonize pancreatic juice than other bacteria. This ability of *Enterococcus* spp. to endure the antibacterial activity of pancreatic juice is, at least in part, due to its higher tolerance of the alkalinity of pancreatic juice than other bacteria.

Previous studies have reported that *E. faecalis* can survive in experimental alkaline condition with pH as high as 11.5^12,13^. This alkaline tolerance has been of interest in particular in the dental field because *E. faecalis* has been isolated from failed endodontic cases, possibly due to its resistance to calcium hydroxide, which has antibacterial activity derived from strong alkalinity^12^. However, pancreatic juice is an alkaline fluid; its pH normally ranges from 8.3 to 8.6 and can be as high as 9.0 due to bicarbonate^14-16^. Therefore, the weak alkalinity of pancreatic juice alone is insufficient to explain the different antibacterial activity against *E. faecalis* compared to that of most other bacteria, which are alkaline sensitive. Nevertheless, *E. faecalis* fitness decreased in pancreatic juice at pH levels of 8.5 or higher in our experiments. This could be due to other antibacterial peptides in pancreatic juice, wherein the pH level is critical for their optimal antibacterial activity^2^. Thus, the mechanism of bacterial colonization in pancreatic juice is hypothesized to contain the following components: (1) triggering events (e.g., slight changes in the pH or antibacterial peptide level of the pancreatic juice, or a change in *E. faecalis* strain) initiate the colonization of *E. faecalis*; (2) *E. faecalis* colonization in pancreatic juice induces further changes to the exocrine function of the pancreas; (3) decreased exocrine function leads to reduced antibacterial activity in pancreatic juice; and (4) these events eventually allow colonization of non-*Enterococcus* bacteria. Indeed, our results also showed enrichment of *Enterococcus* spp. in samples from patients with duodenal tumors, which are supposed to lack chronic pancreatic inflammation. However, the pancreatic juice of patients with pancreatic cancer and bile duct cancer, which may be accompanied by chronic pancreatic inflammation^17^, showed a highly heterogeneous bacterial composition. Clinically, stenting for obstructive jaundice could be an additional factor that promotes heterogeneous bacterial colonization.

Recently, more studies have investigated the effect of the microbiota on cancer^8,18^. Patients with pancreatic cancer exhibit an abundant intratumoral microbiome; some species can induce immunosuppression to promote the progression of pancreatic ductal adenocarcinoma (PDAC)^19^. Additionally, the fungal microbiome can promote pancreatic oncogenesis by driving the complement cascade through the activation of mannose-binding lectin^20^. Bacteria can influence not only the carcinogenesis and cancer progression but also the therapeutic response.

Gammaproteobacteria might contribute to gemcitabine resistance in PDAC^21^. Although the effect of *E. faecalis* on pancreatic cancer is not clear, a possible effect on colorectal cancer can be seen in several reports. However, whether it is tumor promoting or protecting is controversial^22^. Our previous report showed that chronic inflammation and fibrosis exist in the adjacent tissue of pancreatic cancer^17^ and that *E. faecalis* resides in pancreatic cancer tissue of clinical samples^5^. These results suggest that chronic pancreatitis, accelerated by infection, may be involved in the pathogenesis of pancreatic cancer. Additionally, as indicated in this present study, *E. faecalis* can initiate translocation of the microbiome to the pancreas by virtue of its higher survivability in pancreatic juice than that of other bacteria. These results warrant additional future investigation.

## Acknowledgments

We thank the next-generation sequencing core facility of the Genome Information Research Center at the Research Institute for Microbial Diseases of Osaka University for the support in DNA sequencing and data analysis.

## Supplemental Digital contents

Supplemental Figures.docx

Supplemental Table 1.xlsx

## Notes

**Conflicts of Interest and Source of Funding:** The authors have no potential conflicts relevant to the manuscript. This study was supported by Princess Takamatsu Cancer Research Fund and a KAKENHI grant (19H03562) from the Japan Society for the Promotion of Science.

### Competing Interest Statement

The authors have declared no competing interest.

