## Supplemental Figure for "*Enterococcus* spp. have higher fitness for survival, in a pH-dependent manner, in pancreatic juice among duodenal bacterial flora"

Supplemental Figures

### Supplemental Fig. S1

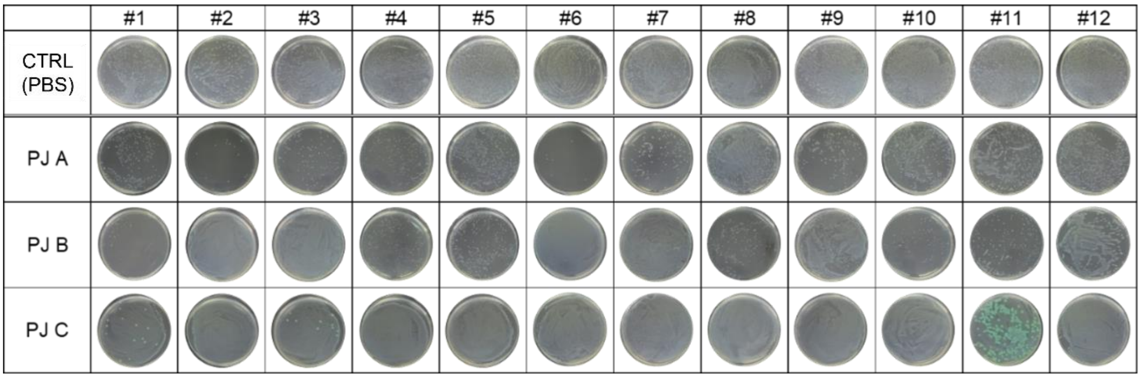


Original images for the visual agar plate assay used in Figure 3A. Twelve isolated E. faecalis (#1-#12) were incubated with each pancreatic juice (PJ A-C) and plated on BHI agar plates. The changes in the bacterial growth compared to control (CTRL) was judged and described in Figure 3A. Unspecified green colony grew in the culture of #1,3 and 11 with PJ C, which was initially characterized as aseptic, suggesting the presence of unspecified bacteria below detectable level in original PJ C. #11 culture in PJ C was judged as non-assessable (NA) due to the contamination, while colonies of #1 and #3 culture in PJ C were clearly increased.

### Supplemental Fig. S2

| score | colony forming unit | |
|---|---|---|
| 0 | **0** | 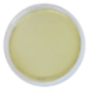 |
| 1 | **0~20** | 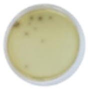 |
| 2 | **21~100** | 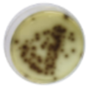 |
| 3 | **More than 100** | 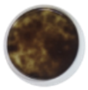 |
| 4 | **uncountable** | 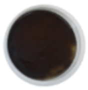 |

A legend for the scoring used in the Figure 3B. Bacterial growth was scored according to the colony number as indicated.

Supplemental Fig. S3

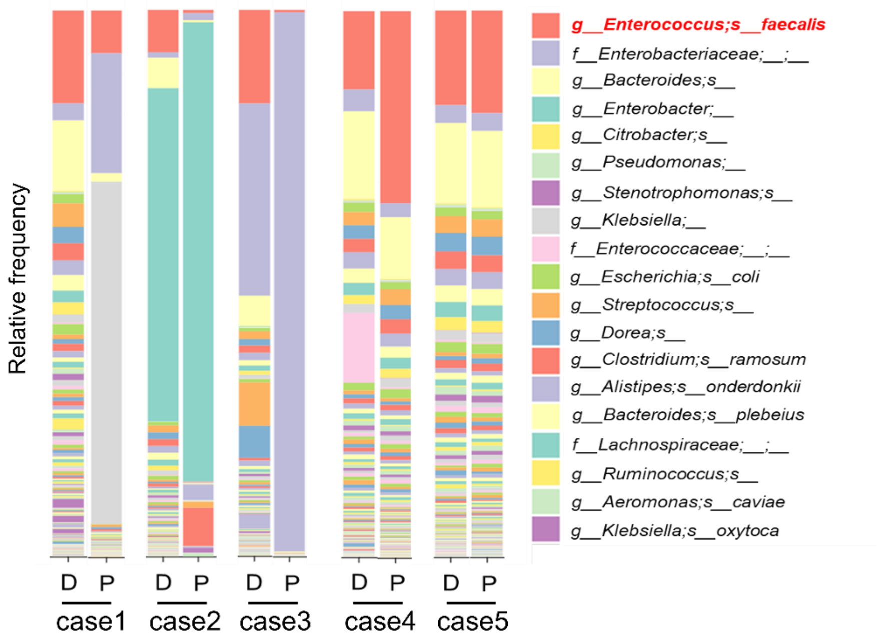


Metagenome analysis of the paired duodenal and pancreatic juice from five pancreatic cancer patients. Each colored bar indicates the relative frequency of indicated bacterial species, which are listed on the right panel. D, duodenal juice; P, pancreatic juice.;
