## Supplemental Table 1 for "*Enterococcus* spp. have higher fitness for survival, in a pH-dependent manner, in pancreatic juice among duodenal bacterial flora"

Supplemental Table1.

Primer list

| PCR specificity | primer name | sequence (5'-3') | PCR conditions |
| --- | --- | --- | --- |
| All bacteria | Universal 16S Fwd | CCTACGGGNGGCWGCAG | 94°C 5min, (94°C 40sec, 55°C 2min, 72°C 1min) 30cycles |
|  | Universal 16S Rev | GACTACHVGGGTATCTAATCC |  |
| *E. faecalis* | Faecalis 16S Fwd | ACACTTGGAAACAGGTGC | 95°C3min, (95°C 30sec, 55°C 30sec, 72°C 25sec) 40cycles |
|  | Faecalis 16S Rev | AGTTACTAACGTCCTTGTTC |  |
